# Fibrillin-Related Proteins Control Calcium Homeostasis in Dystrophic Muscle Across Species

**DOI:** 10.1101/2025.06.09.658655

**Authors:** D Marchiafava, AG Vidal-Gadea

## Abstract

Duchenne muscular dystrophy (DMD) involves progressive muscle degeneration associated with calcium dysregulation, but the mechanisms linking extracellular matrix (ECM) integrity to calcium homeostasis remain unclear. We investigated whether MUA- 3, a fibrillin-related ECM protein in *Caenorhabditis elegans*, contributes to calcium regulation in dystrophic muscle. Using fluorescent calcium imaging in transgenic worms expressing muscle-specific GCaMP2, we found that downregulating *mua-3* selectively elevated resting calcium levels in healthy muscle but had no effect in dystrophic (*dys-1*) muscle, suggesting impaired MUA-3 function in dystrophy. Despite altered calcium dynamics, *mua-3* downregulation did not affect locomotor function. In human dystrophic myoblasts, we observed significantly elevated sarcoplasmic calcium levels concurrent with substantial downregulation of fibrillin genes FBN1/FBN2. These findings demonstrate that fibrillin-related proteins regulate calcium homeostasis across species, suggesting that ECM integrity directly contributes to cellular calcium control in muscle. This work identifies a conserved mechanism linking extracellular matrix stability to intracellular calcium regulation and suggests that targeting ECM-calcium coupling may offer new therapeutic approaches for muscular dystrophy.

## Description

Duchenne Muscular Dystrophy (DMD) is a severe X-linked neuromuscular disorder characterized by progressive muscle degeneration and weakness. DMD results from loss-of-function mutations in the dystrophin (*DMD*) gene, leading to the absence of dystrophin, a critical cytoskeletal protein that connects the intracellular actin cytoskeleton to the extracellular matrix via the dystrophin-associated protein complex (DAPC) (Duan et al., n.d.). The loss of dystrophin leads to the destabilization of the DAPC and compromises the structural integrity of the sarcolemma, rendering it susceptible to damage during muscle contraction and stretching (Allen et al., 2016). Consequently, repeated mechanical stress leads to sarcolemmal tears, increased sarcoplasmic calcium influx, proteolytic enzyme activation, and eventual muscle fiber necrosis (Mareedu et al., 2021). Over time, the regenerative capacity of muscle tissue diminishes, resulting in replacement of muscle fibers with fibrotic and adipose tissue.

In the nematode *Caenorhabditis elegans*, the gene MUA-3 encodes a transmembrane cell adhesion receptor that shares functional properties with human fibrillin-1 and 2 (FBN1, FBN2), which are components of connective tissue microfibrils and the extracellular matrix (ECM)(Davis & Summers, 2012; Ramirez & Sakai, 2010). MUA-3 localizes to hypodermal cells, facilitating the attachment of body wall muscles to the overlying cuticle, and is essential for maintaining the integrity of hemidesmosome-like structures that anchor muscle cells in the worm (Bercher et al., 2001; Fotopoulos et al., 2015). Mutations in *mua-3* lead to progressive muscle detachment during larval development into adulthood, highlighting its critical role in muscle adhesion and structural stability. Given that both MUA-3 and human fibrillins are essential for ECM integrity and tissue mechanical properties, we hypothesized that dysfunction of these fibrillin-related proteins contributes to altered calcium homeostasis in dystrophic muscle. To investigate this, we used a synergistic approach of worm and human myocyte cultures to investigate the role of these proteins in the pathology of DMD.

To understand the role of MUA-3 in maintaining muscle calcium homeostasis, we downregulated *mua-3* in both healthy (ZW495) and dystrophic (AVG6) worms expressing muscle-specific GCaMP protein, using fluorescence as an indirect indicator of calcium (Chen et al., 2013; Nakai et al., 2001). Downregulating *mua-3* did not increase calcium levels during contraction in either wild-type or *dys-1* worms (**Fig. 1B**). However, downregulating *mua-3* selectively increased calcium levels only in the relaxed muscles of healthy worms (**Fig. 1C**). Downregulating *mua-3* had no effect on the relaxed muscle brightness in dystrophic worms. Downregulating *mua-3* also decreased the contracted:relaxed calcium ratio in healthy but not dystrophic worms (**Fig. 1D**). The lack of a calcium increase when downregulating *mua-3* in dystrophic worms suggests that *mua-3* may not function properly in dystrophic worms.

**Figure 1.**
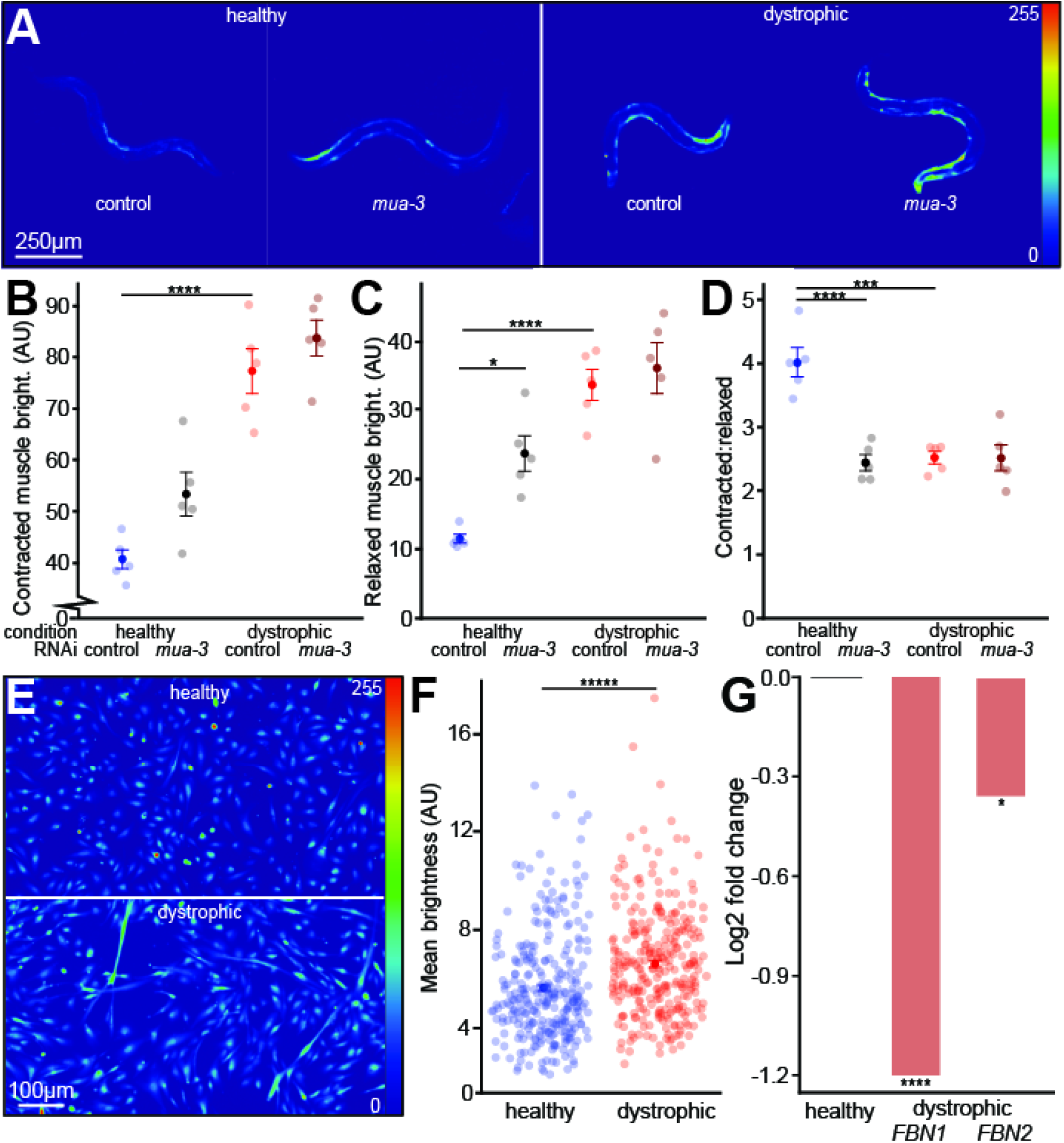
MUA-3 and FBN1/2 contribute to elevated baseline calcium in muscle. **A)** Representative images of brightness levels of wildtype and dystrophic strains in both the control and RNAi treated conditions via *Pmyo3::gCaMP* expression. Brightness levels were linearly adjusted for each image to enhance the visualization of each worm. **B)** Muscle brightness changes in the contracted muscle of each worm (Tukey’s Honest Significance Test, n=50) **C)** Muscle brightness changes in the relaxed muscle of each worm (Tukey’s Honest Significance Test, n=50) **D)** Contracted:relaxed muscle brightness ratio changes of each worm (Tukey’s Honest Significance Test, n=50) **E)** Representative images depicting brightness levels of healthy and dystrophic myoblasts via Fluo-4 AM. Brightness levels were linearly adjusted for each image to enhance visualization. **F)** Dystrophic myoblasts contain elevated sarcoplasmic calcium compared to healthy myoblasts after 1 day of differentiation (unpaired two-tailed t-test, n=300) **G)** Fold change differences in *FBN1* and *FBN2* expression in dystrophic compared to healthy myoblasts via RNASeq of healthy and dystrophic myoblasts (Wald test, n=6). *=p<0.05, **=p<0.01, **\*\***\*=p<0.001 ****=p<0.0001, *****=p< 0.00000001, ******=p<1.88E-13.

Next, we subjected worms to bouts of swimming and recorded the swimming frequency at both the beginning of the bout and after 3 h of swimming in liquid NGM media to measure exercise-induced fatigue. While healthy and dystrophic worms treated with an L4440 empty vector control significantly differed in swimming frequency at both the pre- and post-exercise periods, no significant differences were observed when *mua-3* was downregulated in each condition. Healthy worms with downregulated *mua-3* were still able to swim at a significantly higher frequency than dystrophic worms (Fig. 1E). These results suggest that MUA-3 is not required for proper swimming in healthy or dystrophic worms. Although downregulating *mua-3* increased resting calcium levels in wild-type worms, they were able to maintain their swimming ability.

Next, we aimed to validate these findings in human myocyte cultures, where FBN1 and FBN2 represent the fibrillin-related proteins analogous to MUA-3 function in ECM integrity. Using Fluo-4 AM calcium indicator dye, we quantified calcium levels in healthy and dystrophic myoblasts after one day of differentiation. Dystrophic myoblasts displayed significantly elevated calcium levels compared to healthy controls (**Fig. 1F/G**). To examine FBN1/FBN2 expression, we analyzed RNASeq data from an independent, unpublished study (Marchiafava et al., in preparation), which revealed significant downregulation of both genes in dystrophic myoblasts.

Collectively, our data indicate that downregulation of FBN1 and FBN2 is associated with elevated resting sarcoplasmic calcium levels in dystrophic human myoblasts, mirroring the calcium dysregulation observed upon downregulating *mua-3* in *C. elegans*. The absence of calcium elevation in contracted muscles of *mua-3* downregulated dystrophic worms suggests that dystrophic conditions may impair MUA-3 function, potentially compromising sarcolemmal integrity. This is further supported by the parallel downregulation of fibrillin-related proteins (FBN1/FBN2) in human dystrophic myoblasts. Given the established role of fibrillins in organizing ECM-integrin-cytoskeleton linkages critical for sarcolemmal stability, we propose that FBN1/FBN2 deficiency could contribute to calcium dyshomeostasis in dystrophic muscle by weakening cell-ECM coupling.

## Methods

### *C. elegans* strains and culture conditions

Animals were maintained at 20°C on nematode growth media (NGM) agar plates seeded with *Escherichia coli* OP50 as previously described (Brenner, 1974). Strains used in this study included the ZW495 strain (*zwIs132[myo-3p::GCaMP2 + lin-15(+)]*)and the AVG6 strain generated by crossing ZW495 with the BZ33 strain (*dys-1(eg33))*. All strains were obtained from the Caenorhabditis Genetics Center (CGC) which is supported by the NIH Office of Research Infrastructure Programs (P40 OD010440).

### Calcium imaging in *C. elegans* embryos

Animals carrying *zwIs132[Pmyo- 3::GCaMP2::unc-54_3’UTR]* were imaged as previously described (Fox et al., 2007; Hughes et al., 2019). Motion recording was followed by single-frame fluorescence imaging under blue-light illumination using a Sola Light Engine. Images were acquired to minimize fluorophore bleaching.

### Calcium quantification in *C. elegans*

ImageJ was used to draw a region of interest (ROI) around each specified region of animal muscle to measure maximum GCaMP2 brightness (indicating relative calcium levels) (Schneider et al., 2012). A matched ROI outside the animal served as background. The brightness ratio was calculated as animal fluorescence in the region of interest divided by adjacent background fluorescence.

### *C. elegans* RNA interference (RNAi)

WT (ZW495) and dystrophic (AVG6) worms were cultivated on NGM plates containing IPTG and seeded with bacteria expressing either the L4440 empty vector (control) or RNAi targeting *mua-3*. Downregulation of gene expression was confirmed via qPCR.

### *C. elegans* swimming assays

Day-one adults were placed in a 12-well agar plate and flooded with 1mL liquid NGM. Animals were filmed and swimming frequency was calculated as the number of body bends per second once placed in the NGM as well as after a 3-hour period to quantify endurance.

### Human myocyte cultures

Age matched (16 yo male) immortalized healthy (AB1190) and dystrophic (Ab1071) skeletal myoblast lines (Myology Institute in France) were maintained in Skeletal Muscle Growth Media (PromoCell) on Matrigel (1%, Fisher, Cat #CB-40234A) until ∼40% confluence. All experiments used cells within six passages.

### Human myocyte calcium imaging

Myoblasts were seeded at 1×10^5^ cells/mL on Matrigel-coated 12-well plates and loaded with 5 μM Fluo 4-AM (Thermo Fisher) in growth medium for 30 minutes at 37°C, then washed and imaged in live imaging medium (Gibco) using the Keyence BZ-X810 fluorescent microscope. ImageJ was used to calculate mean brightness of each myoblast.

### RNASeq

Unpublished RNASeq data (Marchiafava et al., *in preparation*) are available from the authors upon reasonable request. RNA was extracted from wildtype or dystrophic myoblasts using the DirectZol RNA Miniprep Kit (Zymo). RNA-seq library preparation and sequencing were performed by Azenta Life Sciences. Briefly, mRNA was enriched using poly-A selection, and libraries were prepared using the NEBNext Ultra II RNA Library Prep Kit for Illumina (NEB). Paired-end sequencing was performed targeting a minimum depth of 30 million reads per sample.

## Acknowledgements

Strains were provided by the Caenorhabditis Genetics Center (funded by NIH grant P40 OD010440).

## Funding

Funding was provided by the National Institutes of Health, National Institute of Arthritis and Musculoskeletal and Skin Diseases award 2R15AR068583-02.

## Author Contributions

Damiano Marchiafava: Conceptualization, Project administration, Methodology, Visualization, Writing original draft.

Andres Vidal-Gadea: Conceptualization, Project administration, Resources, Funding acquisition, Supervision, Writing (review & editing).

## Conflict of Interest

The authors declare no conflict of interest.

## Notes

### Competing Interest Statement

The authors have declared no competing interest.

## References

Allen, D. G., Whitehead, N. P., & Froehner, S. C. (2016). Absence of Dystrophin Disrupts Skeletal Muscle Signaling: Roles of Ca2+, Reactive Oxygen Species, and Nitric Oxide in the Development of Muscular Dystrophy. Physiological Reviews, 96(1), 253–305. 10.1152/physrev.00007.2015

Bercher, M., Wahl, J., Vogel, B. E., Lu, C., Hedgecock, E. M., Hall, D. H., & Plenefisch, J. D. (2001). Mua-3, a gene required for mechanical tissue integrity in Caenorhabditis elegans, encodes a novel transmembrane protein of epithelial attachment complexes. The Journal of Cell Biology, 154(2), 415–426. 10.1083/jcb.200103035

Brenner, S. (1974). THE GENETICS OF CAENORHABDITIS ELEGANS. Genetics, 77(1), 71–94. 10.1093/genetics/77.1.71

Chen, T.-W., Wardill, T. J., Sun, Y., Pulver, S. R., Renninger, S. L., Baohan, A., Schreiter, E. R., Kerr, R. A., Orger, M. B., Jayaraman, V., Looger, L. L., Svoboda, K., & Kim, D. S. (2013). Ultrasensitive fluorescent proteins for imaging neuronal activity. Nature, 499(7458), 295–300. 10.1038/nature12354

Davis, M. R., & Summers, K. M. (2012). Structure and function of the mammalian fibrillin gene family: Implications for human connective tissue diseases. Molecular Genetics and Metabolism, 107(4), 635–647. 10.1016/j.ymgme.2012.07.023

Duan, D., Goemans, N., Takeda, S., Mercuri, E., & Aartsma-Rus, A. Duchenne muscular dystrophy. Nature Reviews Disease Primers, 7(1), 13.

Fotopoulos, P., Kim, J., Hyun, M., Qamari, W., Lee, I., & You, Y.-J. (2015). DPY-17 and MUA-3 Interact for Connective Tissue-Like Tissue Integrity in Caenorhabditis elegans: A Model for Marfan Syndrome. G3 Genes|Genomes|Genetics, 5(7), 1371–1378. 10.1534/g3.115.018465

Fox, R. M., Watson, J. D., Von Stetina, S. E., McDermott, J., Brodigan, T. M., Fukushige, T., Krause, M., & Miller, D. M. (2007). The embryonic muscle transcriptome of Caenorhabditis elegans. Genome Biology, 8(9), R188. 10.1186/gb-2007-8-9-r188

Hughes, K. J., Rodriguez, A., Flatt, K. M., Ray, S., Schuler, A., Rodemoyer, B., Veerappan, V., Cuciarone, K., Kullman, A., Lim, C., Gutta, N., Vemuri, S., Andriulis, V., Niswonger, D., Barickman, L., Stein, W., Singhvi, A., Schroeder, N. E., & Vidal-Gadea, A. G. (2019). Physical exertion exacerbates decline in the musculature of an animal model of Duchenne muscular dystrophy. Proceedings of the National Academy of Sciences, 116(9), 3508–3517. 10.1073/pnas.1811379116

Mareedu, S., Million, E. D., Duan, D., & Babu, G. J. (2021). Abnormal Calcium Handling in Duchenne Muscular Dystrophy: Mechanisms and Potential Therapies. Frontiers in Physiology, 12, 647010. 10.3389/fphys.2021.647010

Nakai, J., Ohkura, M., & Imoto, K. (2001). A high signal-to-noise Ca2+ probe composed of a single green fluorescent protein. Nature Biotechnology, 19(2), 137–141. 10.1038/84397

Ramirez, F., & Sakai, L. Y. (2010). Biogenesis and function of fibrillin assemblies. Cell and Tissue Research, 339(1), 71–82. 10.1007/s00441-009-0822-x

Schneider, C. A., Rasband, W. S., & Eliceiri, K. W. (2012). NIH Image to ImageJ: 25 years of image analysis. Nature Methods, 9(7), 671–675. 10.1038/nmeth.2089

